# Selective bacteriophages reduce the emergence of resistant bacteria in the bacteriophage-antibiotic combination therapy

**DOI:** 10.1101/2023.01.22.525106

**Authors:** Aa Haeruman Azam, Koji Sato, Kazuhiko Miyanaga, Tomohiro Nakamura, Shinjiro Ojima, Kohei Kondo, Azumi Tamura, Wakana Yamashita, Yasunori Tanji, Kotaro Kiga

## Abstract

*Escherichia coli* O157:H7 is a globally important foodborne pathogen that affects food safety. Antibiotic administration against O157:H7 may contribute to the exacerbation of hemolytic uremic syndrome (HUS) and antibiotic-resistant strains increase; therefore, bacteriophage therapy (phage therapy) is considered a useful alternative. In the treatment of resistant bacterial infections, combination therapy with bacteriophages and antibiotics, taking advantage of the benefits of both agents, has been suggested to be effective in inhibiting the emergence of antimicrobial-resistant strains; however, its effectiveness against O157:H7 is not well understood. In this study, we isolated SP015, a phage that infects O157:H7, and compared the combined effect of the bacteriophage and fosfomycin (FOM) with that of the PP01 phage. Genomic analysis revealed that FOM exerts its antibacterial activity through glycerol-3-phosphate transporter (GlpT) and hexose phosphate transporter (UhpT) proteins, and the receptors of PP01 and SP015 phages are the outer membrane protein C (OmpC) and ferrichrome outer membrane transporter protein (FhuA), respectively. Experiments with knockout strains have suggested that FOM also uses OmpC, the receptor for PP01, as a transporter. This may explain why the combination treatment with PP01 resulted in a faster emergence of resistance than the combination treatment with SP015. We propose that phage-antibiotic combination therapy requires careful selection of the phage to be used.

## Introduction

Antibiotics have been used in clinical practice to fight infectious diseases since their discovery about a century ago (1). However, the extensive use of antibiotics and lack of effective control treatments have led to a rapid increase in the emergence of antimicrobial-resistant (AMR) bacteria. It is projected that by 2050, the number of deaths caused by AMR infection will outnumber the deaths caused by cancer if alternative treatments are not available (2). The use of bacteriophages, viruses that specifically infect bacteria (phage therapy), has attracted significant attention as a possible alternative to combat the AMR problem, and phage therapy has been used in some countries (3–11). However, the emergence of phage-resistant bacteria, including clinical experiments in humans, has been reported in several studies (12–14). Therefore, rather than replacing antibiotics with phages, combination therapy is being used to combat AMR. Combination therapy is known to be less likely to produce resistance than either therapy alone.

Enterohemorrhagic *Escherichia coli* serogroup O157:H7 is a worldwide source of infection that causes bloody diarrhea and hemolytic uremic syndrome (HUS) in humans and animals (15–17). Most *E. coli* O157:H7 infections in humans are foodborne diseases that are transmitted through domestic animals as reservoirs of O157:H7 (17, 18). The use of phages to control pathogenic organisms in the gastrointestinal tract is promising, and several studies have been successful in animal models (19–22).

Fosfomycin (FOM), an antibiotic produced by *Streptomyces* sp. in 1969, has been used for many years (23). As the number and types of drug-resistant bacteria are increasing, FOM is currently attracting attention as an effective antibacterial drug against multidrug-resistant bacteria owing to its low molecular weight (molecular weight:138) compared to other antibiotics (24, 25). Currently, FOM is used as a standard treatment for urinary tract infections caused by *E. coli* and fecal streptococci, and hemolytic uremic syndrome caused by enterohemorrhagic *E. coli* (26, 27). Studies have also reported the efficacy of FOM against multidrug-resistant *E. coli* and *Pseudomonas aeruginosa* (28, 29). However, FOM alone may increase the probability of developing HUS and should be used with caution in the clinical setting.

For both antibiotics and phages, tradeoffs occur with the acquisition of resistance. In both cases, changes in the membrane structure and reduced growth rates frequently occur; however, the mechanisms of phage resistance and antibiotic resistance are usually different. This means that a combination of antibiotics and phages administered simultaneously may be more effective than administration alone because different trade-offs are required for the bacteria. Although simultaneous administration of antibiotics and phages may be effective against O157:H7, its efficacy has not yet been investigated. Therefore, our aim was to evaluate the potential use of two different phages, PP01 and SP015, that infect O157:H7 and analyze the underlying resistance mechanism of the bacteria exposed to the phages and FOM combined or to each antimicrobial alone.

## Material and methods

### Media and buffer

All experiments were conducted using Mueller-Hinton broth (2 g beef extract, 17.5 g acid digest casein, 1.5 g soluble starch per liter) and Luria Bertani (LB) broth (10 g polypeptone, 10 g sodium chloride, and 5 g yeast extract per liter). In accordance with the method used for testing drug susceptibility to FOM, glucose-6-phosphate (G6P) was added after autoclaving the medium to a final concentration of 25 μg/mL (30). Hereafter, this medium will be referred to as MHB-G6P. Fosfomycin was dissolved in sterile water, sterilized through a 0.22 μm filter, and stored at -20°C.

Phosphate-Buffered Saline (PBS) (8.0 g NaCl, 0.2 g KCl, 1.44 g Na_2_HPO_4_, and 0.24 g KH_2_PO_4_) was used for dilution of the bacteria solution and Sodium-Magnesium (SM) buffer (5.8 g NaCl, 0.2 g MgSO_4_.7H_2_O, 50 ml 1M Tris-HCl (pH. 7.5), and 5 mL of 2% (w/v) Gelatin solution were used for phage solution dilution.

### Bacteria and Phages

The strains and phages used in these experiments are listed in Supplementary Table S1. *E. coli* O157:H7 ATCC 43888 (hereafter O157) has the same serotype as pathogenic *E. coli* O157:H7, but is nonpathogenic as it does not possess the Shiga toxin genes, *stx1* and *stx2* (31). PP01 and SP015 are lytic phages that can infect several O157: O157 strains; hence, the phages were propagated using O157 as a host. Phage solutions were prepared using the overlaid agar plate method, as previously described (32–34). Briefly, 100 μL each of 10^5^–10^6^ PFU/mL (PFU: Plaque Forming Unit) phage solution and O157 pre-culture solution were mixed, and the mixture was added to 3 mL soft agar dissolved at 45°C, layered on LB plates, and incubated at 37°C overnight. Afterwards, 4 mL of SM buffer was added to the plate on which the plaque was formed, only the upper layer was scraped off, and the supernatant was collected after centrifugation (10,000 ×g, 5 min, 4°C). Finally, chloroform was added to a final concentration of 2% (v/v) to break up the remaining phage, which was stored at 4°C. Phage concentration was measured using the plaque assay method. Diluted phage solution (10^3^–10^4^ PFU/mL) and O157 pre-culture solution were mixed in 100 μL portions, added to 3 ml soft agar dissolved at 45°C, layered on an LB plate, and incubated at 37°C overnight, and the phage solution concentration was determined by counting the number of plaques.

### Isolation of resistant strains and mutant phage

Isolation was performed for each stock obtained from the passage co-culture. For resistant O157 strains, a portion of each glycerol stock was streaked on an MHB-G6P plate, incubated overnight at 37°C. Afterwards, one single colony was taken, inoculated into 2 mL of MHB-G6P, and incubated overnight at 37°C, 120 rpm. The mutant phage was propagated using wild-type *E. coli* O157 in LB medium.

### Isolation and preparation of phage stock

Phage SP015 was isolated from sewage influent obtained from a municipal wastewater treatment plant in Tokyo using *E. coli* O157:H7 as the propagation host using the double-layer agar plating method. The phages were propagated and purified using a previously described method(35). Briefly, the purified phage was propagated by mixing 1% of the overnight culture of O157 in liquid LB and incubated overnight at 37°C. Host cells were removed through centrifugation (11,000 ×g, 20 min, 4°C) before phage concentration using the polyethylene glycol 6000-NaCl (PEG-NaCl) method and filtered through a 0.22 μm Millex-GP filter (Merck, Millipore, Darmstadt, Germany).

### TEM imaging of phages

Phages were observed using transmission electron microscopy (TEM), as described previously (36). Briefly, PEG-NaCl concentrated phage lysate was purified through cesium chloride (CsCl) step centrifugation (step densities, 1.46, 1.55 and 1.63 g/mL), and concentrated phage suspension 10^9^ plaque forming unit (PFU/mL) were spotted on top of a hydrophilic plastic-carbon-coated copper grid (Nissin EM Corporation, Tokyo, Japan). Phages were allowed to adsorb for 1 min before removing the excess samples. Subsequently, 10 mL of distilled water was spotted onto the grid and removed after a short time. Phages were stained with 2% uranyl acetate or an EM Stainer (Nissin EM Corporation). Excess stain was removed after 1 min, and the grid was allowed to air-dry for 30 min before observation using a JEOL JEM-1400Plus (TEM) operating at 80 kV.

### Characterization of phage growth and determination of phage host range

A one-step growth curve was constructed to determine the burst size and latent period as previously described, with some modifications(36). Briefly, the phages were added to a refreshed overnight culture of bacteria (OD_660_ = 1) at a multiplicity of infection (MOI) of 0.01, and incubated at 37 °C for 10 min, with shaking at 120 rpm. The unbound phages were removed through centrifugation and washed five times with chilled LB medium. Phage-infected cells were incubated at 37 °C for 1 h. The number of phages was determined by the double-layer agar method. The host range was determined using 17 strains of *Escherichia coli* (See Table 1) by dropping 2.5 μL of phage lysate of 10^7^ PFU/mL on bacteria mixed with 0.5% (w/v) top agar.

### Evaluation of phage infectivity through spot test

To evaluate the infectivity of phage-resistant and mutant phages obtained from each culture, a spot test was performed:100 μL of the overnight culture of O157 (wild-type or resistant clone) was added to 3 ml LB soft agar, poured on LB plates, and allowed to dry for 10 min. Then, 5 μL of 10^9^ PFU/mL phage (wild-type or mutant phage) was dropped onto the plate and allowed to stand until it dried. After overnight incubation at 37°C, formation of inhibition circles (plaques) was recorded.

### Phage adsorption assay

The adsorption efficiency of phages on O157 was measured through titrating the free phages present in the supernatant after 20 min of cell-phage contact at an MOI of 0.01. One hundred microliters of the cell-phage solution was sampled and immediately added to 9.9 mL of chilled SM buffer. The solution was gently vortexed before taking 1 mL for centrifugation (10,000 ×g, 5 min, 4 °C) to remove the bacterial cells before titrating the phage concentration. Adsorption efficiency was calculated by dividing the number of adsorbed phages by the initial number of phages. Statistical analysis was performed using two-tailed Student’s t-test.

### Minimum inhibitory concentration (MIC) measurement test

To determine the susceptibility of O157 to FOM, a minimum inhibitory concentration (MIC) assay test was performed. The measurements were performed according to the method prescribed by the CLSI (30). Briefly, the O157 culture medium was diluted in PBS to 10^8^ CFU/mL (CFU: colony-forming unit). Subsequently, MHB-G6P plates containing FOM at 2-fold diluted concentrations (0, 1, 2, 4, … [μg/mL]) were prepared and 1μL of the diluted culture medium was dropped onto the plates. After drying, the plates were incubated at 35°C for 16-20 hours, and the MIC was defined as the minimum concentration of FOM at which no colonies grew.

### Acquisition of antibiotic and phage-resistant strains

To screen for antibiotic- and phage-resistant strains, O157 was passaged in medium containing FOM and phage (PP01 or SP015). L-shaped test tubes containing 4 mL of MHB-G6P were inoculated with 10^7^ CFU/mL of O157 overnight culture, and FOM (4 μg/mL) and PP01 or SP015 (10^7^ PFU/mL) were added alone or in combination 1 h after bacterial addition. Those to which neither FOM nor phage was added were used as controls. Cultures were incubated using a small shaking culture device (Biophotorecorder TVS062CA, ADVANTEC) at 37°C with shaking at 40 rpm, and the turbidity (OD_660_) was measured every 15 min. The incubation time was allowed to run until the bacteria growth reached a stationary phase, which was 48 h when used alone, and 72 h when used in combination. After the end of incubation, the phage concentration in the culture medium was measured using the plaque assay method for those to which phages were added. In the following round, 1% inoculation in 4 mL of new MHB-G6P was performed from the previous round of control and those to which only phage was added and cultured again under the same conditions. For those to which FOM was added, the MIC was measured at the end of each round, and FOM was added to 4 mL of fresh MHB-G6P to reach the same concentration as the observed MIC, and then incubated again for the next round. This procedure was repeated five rounds. The culture medium was centrifuged (10000 ×g, 5 min, 4°C) and the supernatant was stored as phage stock, whereas the pellet was stored as bacterial stock in 15% glycerol at 4°C and -60°C, respectively. The same experiment was performed three times (three runs) with phage-only addition (PP01, SP015), and five times (five runs) with FOM addition (FOM alone, PP01+FOM, and SP15+FOM).

### Determination of specific growth rate of resistant strains

The specific growth rate in the absence of antibiotics and phages was determined to clarify the fitness cost of resistant strains obtained under each condition. Four milliliters of L-shaped test tubes containing 4 mL of LB were inoculated with wild-type O157 or resistant strain at 10^7^ CFU/mL and incubated at 37°C and 40 rpm for 12 h. Turbidity (OD_660_) was measured every 15 min during incubation, and the specific growth rate was determined when the OD_660_ value ranged from 0.5 to 1.1, which was considered to be the growth log phase.

### Genome extraction and whole-genome analysis of resistant strains and mutant phages

The GenElute™ bacterial genomic DNA purification kit (Sigma-Aldrich, USA) was used for bacterial genome extraction, and the phage DNA extraction kit (Norgen Biotex Corp.) was used for phage genome extraction, according to the manufacturer’s protocol. Whole-genome sequencing was performed using the Whole Genome Analysis Service of BGI Japan, Inc. The whole-genome sequence data of wild-type O157 (ATCC 43888) were registered with the National Center for Biotechnology Information (NCBI) (Accession number: CP041623). The genome sequence data were compared to those of the same strain in our laboratory. Therefore, we assembled the sequence data of the wild-type and resistant mutant strains held in our laboratory using BWA (ver. 0.7.17), Samtools (ver. 0.1.19), and Pilon (ver. 1.23), with strains registered in the NCBI database as references(37). Based on this assembled wild-type strain data, mapping was performed using the above software to identify mutations present in the resistant strains. Both wild-type PP01 and SP015 were identified for mutations using data from NCBI (PP01: LC348379, SP015: AP019559) as a reference.

### Genetic manipulation in O157

Primers used in this study are listed in Supplementary Table S2. To delete each gene (*uhpT, glpT, ompC*, and *fhuA*) from wild-type O157, recombinant plasmids were constructed using the primers listed in Supplementary Table S1. Recombination templates were prepared using overlap extension PCR to generate the deletion template, as described in Supplementary Fig. S5.

Plasmid pKOV (Addgene, USA) was used to delete the gene of interest, following the established protocol (38). Plasmids and recombination templates were treated with the restriction enzymes BamHI and SalI (New England Biolabs, USA) and ligated with T4 ligase (New England Biolabs, USA). The ligation product was introduced into *E. coli* JM109 using the heat shock method and cultured on a chloramphenicol LB plate. Homologous recombination of the target genes in the O157 genome using pKOV was performed following an established protocol with some modifications (38). Using electroporation (1.8 kV, 25 μF, and 200 Ω), the plasmid was electroporated into O157 cells, plated on a chloramphenicol LB plate, and incubated at 30°C overnight. One to three of the obtained colonies were suspended in 1 mL PBS, streaked on C chloramphenicol LB plates, and incubated at 43°C overnight to allow for plasmid integration. To increase the transfection efficiency, incubation at 43°C was performed twice. Three colonies were selected from the plates, suspended in 1 mL of PBS, plated on LB plates containing 10% (w/v) sucrose, and incubated at 30°C overnight (double crossover). The resulting recombinants were confirmed to be deficient in the target gene using Sanger sequencing (Biotechnology Division, Department of Biotechnology, Tokyo Institute of Technology). To delete multiple genes from a single strain, the above method was performed sequentially, using the corresponding plasmid for each target gene.

## Result

### Phage isolation and characterization

A novel bacteriophage, SP015, was isolated from a wastewater treatment plant in Tokyo, using *E. coli* O157:H7 as a propagation host. We tested the host range of SP015 and compared it with that of our previously isolated phage PP01, a Myovirus phage belonging to Tequatrovirus(39). Unlike PP01, which superficially infect O157:H7, SP015 exhibited broad host range infecting various strains of *E. coli* (Fig. 1A). Morphological and genomic analyses and several physiological tests were performed to identify and characterize SP015. This bacteriophage belongs to the siphovirus group based on the morphology observed, namely a capsid head connected to a long noncontractile tail (Supplementary Fig. S3A). The latent period and burst size of the phage were 20 min and 53 pfu/cell, respectively (Supplementary Fig. S3B). Because adsorption of phages through bacterial receptors is the first important step in phage infection, we attempted to identify the receptor of SP015 by selecting a spontaneous mutant that completely lost susceptibility to SP015. We found that the adsorption of SP015 was significantly impaired in O157 with truncated FhuA and complementation of FhuA *in trans* rescued phage adsorption (Supplementary Fig. S3C), suggesting that FhuA is a receptor of SP015.

**Figure 1.**
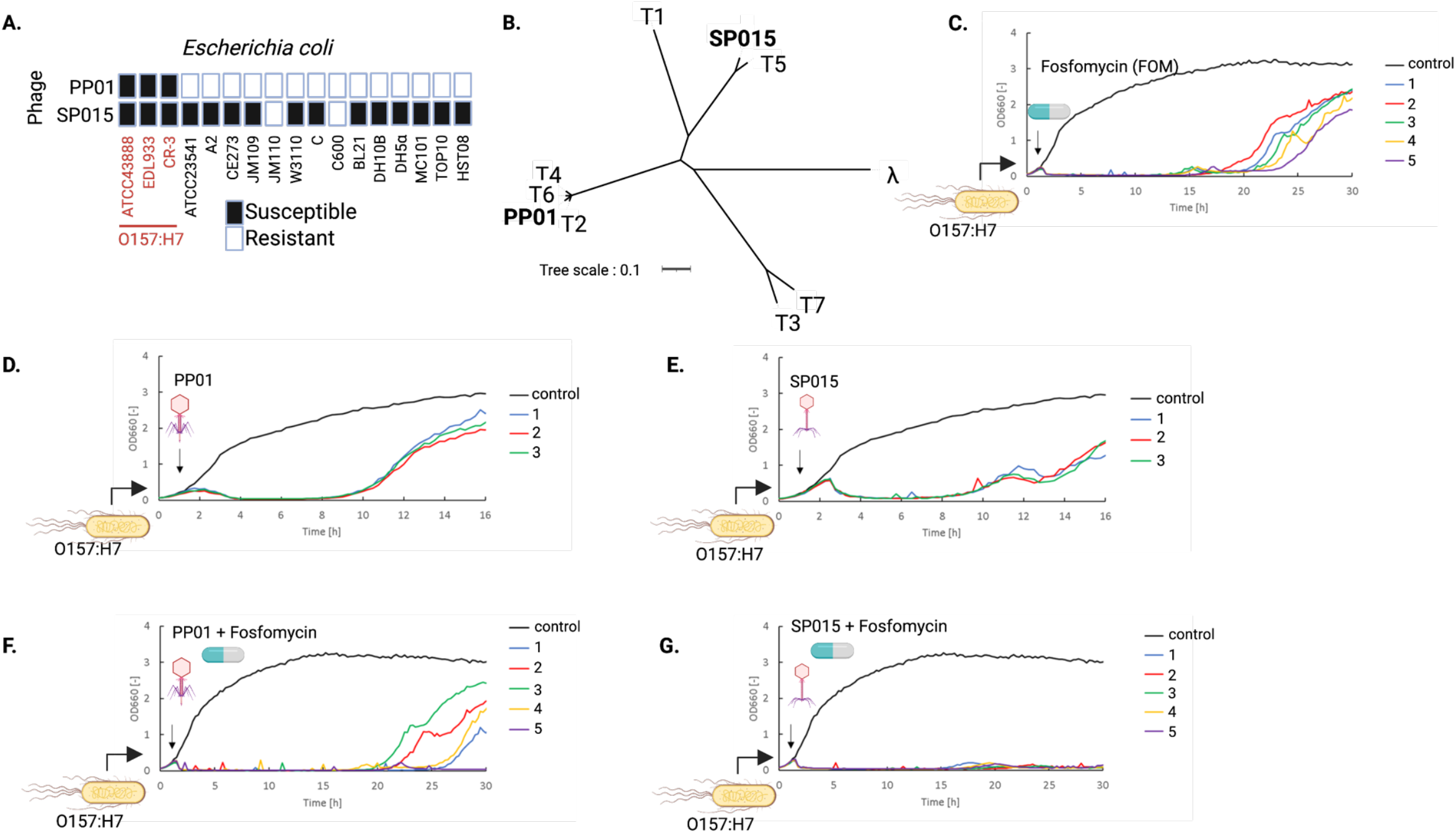
Phage and antibiotic combinations reduces the emergence of resistant strains of O157 more than either treatment alone. (A) Host range of PP01 and SP015 phages against various *Escherichia coli*. (B) Phylogenetic tree of PP01 and SP015 among T-series and lambda phages. (C) Bacterial lysis curve under Fosfomycin (FOM) treatment. FOM with final concentration 4μg/ml was added into culture 1h after bacterial addition. The same experiment was performed in five different runs. (D) Bacterial lysis curve under phage PP01 treatment. The same experiment was performed in three different runs. (E) Bacterial lysis curve under phage SP015 treatment. The same experiment was performed in three different runs. (F) Bacterial lysis under FOM and phage PP01 treatment. The same experiment was performed in five different runs. (G) Bacterial lysis under FOM and SP015 treatment. The same experiment was performed in five different runs. Phage at multiplicity of infection (MOI) =1 or FOM at a final concentration 4 μg/ml were added 1h after bacterial addition.

The assembled SP015 genome consisted of 110,964 bp with 39% G + C content, 163 open reading frames (ORFs), and 23 tRNAs. The phage could be classified as a T5-like coliphage (Fig. 1B) under Tequintavirus, with 90.45% genome identity to T5 (genome accession number: AY543070). Owing to the absence of sequences encoding integrase, recombinase, repressors, and excisionase in its genome, the SP015 phage can be considered a lytic virus. Furthermore, SP015 had no virulence factors or antibiotic resistance genes in its genome, confirming that it is a genetically safe phage suitable for phage therapy.

### The combination of phage SP015 and fosfomycin suppressed the development of resistance

To evaluate the combination therapy of phages and antibiotics, the bacteriophages SP015 and PP01, which infect O157, and Fosfomycin (FOM), a commonly used antibiotic against O157, were examined for their effectiveness in killing O157. We observed that treatment with FOM alone could effectively lyse O157 and suppress the growth of resistant clones until approximately 20 hours (Fig. 1C) whereas phage treatment with either PP01 and SP015 was only effective for less than 20 hours (Figs. 1D and 1E). The addition of PP01 or SP015 one hour after bacterial addition resulted in a decrease in turbidity due to bacterial cell lysis in three independent runs; however, approximately eight hours later, turbidity increased due to the appearance and growth of phage-resistant bacteria (Figs. 1D and 1E). With SP015 addition, in all runs, the turbidity decreased again at approximately 11.5-hour point but increased at 13.5-hour (Fig. 1E).

However, combination treatment of phage and FOM displayed different outcomes depending on the type of phage used; combination treatment of PP01 and FOM showed an extension of the growth inhibition time in run 5 but similar effect as of FOM alone in the other four runs (Fig. 1F), on the other hand, combination treatment of SP015 and FOM could delay the emergence of resistant clone in all runs for up to 30 hours (Fig. 1G).

### Combination use of phage and fosfomycin delayed the occurrence of fosfomycin-resistant O157

To study the arm-race of phages and bacteria during their interaction with or without fosfomycin, we continued the incubation of the bacteria shown in Fig. 1 for five rounds, as depicted in Fig. 2A. In the co-culture of PP01 and O157, bacterial lysis was observed during the first round in all three independent runs, and was observed again in the third round of run 3 (Fig. 2B). In contrast, bacterial lysis occurred only in the first round of all runs of co-culture between SP015 and O157 (Fig. 2C).

**Figure 2.**
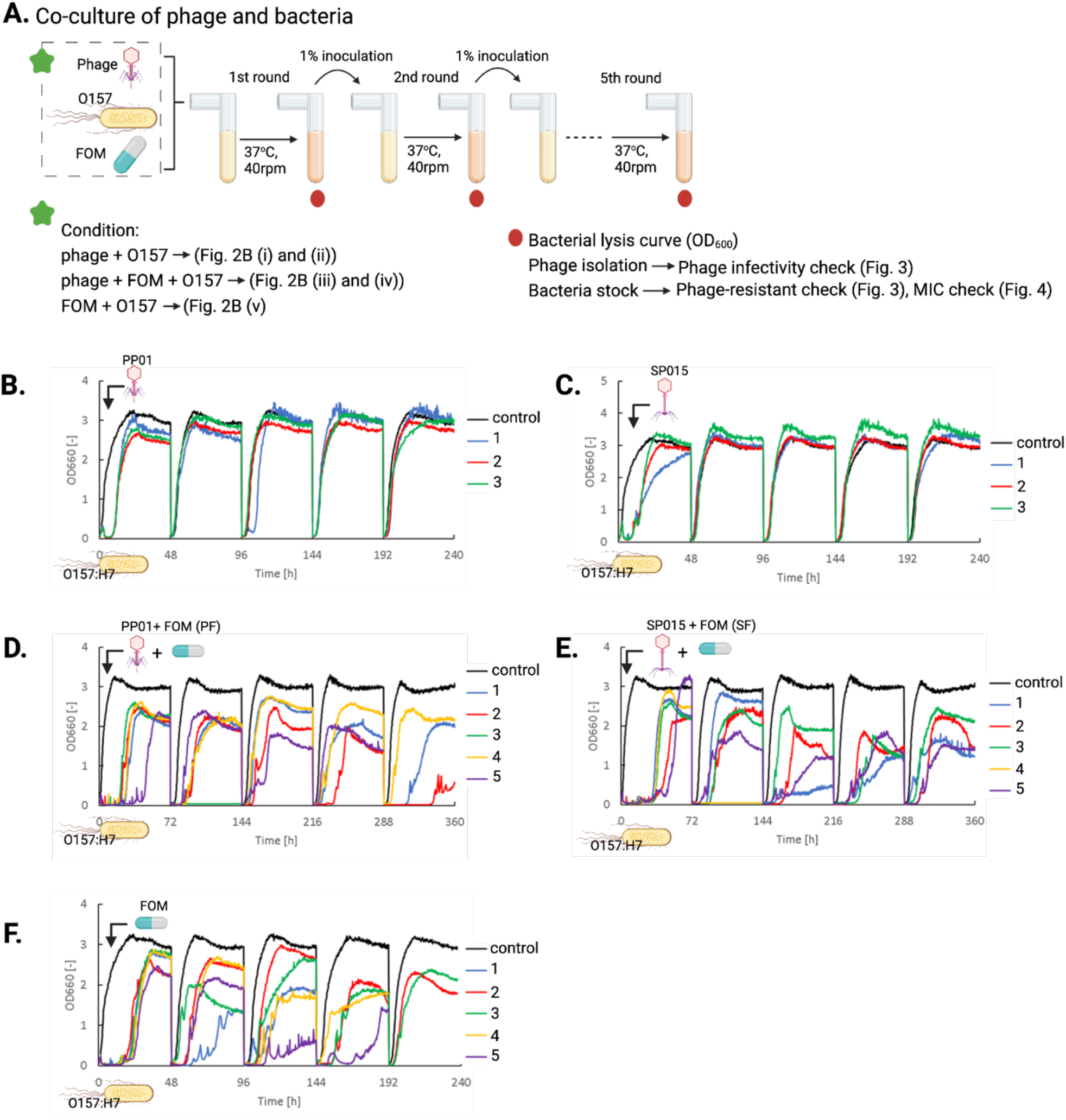
Phage quantification, MIC test, and genomic analysis of resistance bacteria from extended passage co-culture of phage and bacteria. (A) Illustration of passage co-culture of phage and bacteria with or without addition of FOM. The co-culture was performed until fifth rounds where phage and bacteria were collected at final passage and subjected for whole genome analysis. (B) Bacterial lysis curve during five rounds of co-culture of bacteria and phage PP01 in the absence of FOM. (C) Bacterial lysis curve during five rounds of co-culture of bacteria and phage SP015 in the absence of FOM. (D) Bacterial lysis curve during five rounds of co-culture of bacteria and phage PP01 in the presence of FOM. (E) Bacterial lysis curve during five rounds of co-culture of bacteria and phage SP015 in the presence of FOM. (F) Bacterial lysis curve during five round of culture with FOM addition.

In the presence of FOM alone or in the presence of phages PP01 or SP015, bacterial lysis was apparent in all rounds (Fig. 2D - F). The growth of resistant bacteria was not observed in run 3 after the second round and in run 5 after the fifth round of PP01+FOM (PF) co-culture, indicating the complete eradication of bacteria (Fig. 2D). Similarly, complete eradication of O157 was observed in run 4 of the SP015+FOM (SF) co-culture (Fig. 2E).

Next, we investigated whether the combination therapy could suppress the emergence of antibiotic resistance. To do so, we determined the MIC values of resistant bacteria isolated from each round of FOM, PF, and SF. We found that the MIC of FOM against wild-type O157 (ATCC 43888) was 16 μg/mL. According to the criteria for FOM resistance established by CLSI, bacteria with MIC values of ≥ 256, 128, and ≤ 64 μg/mL were considered to be resistant, intermediate-resistant, and sensitive bacteria, respectively. Therefore, the wild-type O157 strain used in this study is sensitive to FOM.

Combination therapy of either phage PP01 or SP015 with FOM significantly delayed the emergence of resistant O157, independent of the difference in the type of phage used (Fig. 3). The average MIC value of resistant bacteria from phage and FOM treatment showed a tendency to have lower MIC than those that were isolated from FOM treatment alone around 2.5 order of magnitude.

**Figure 3.**
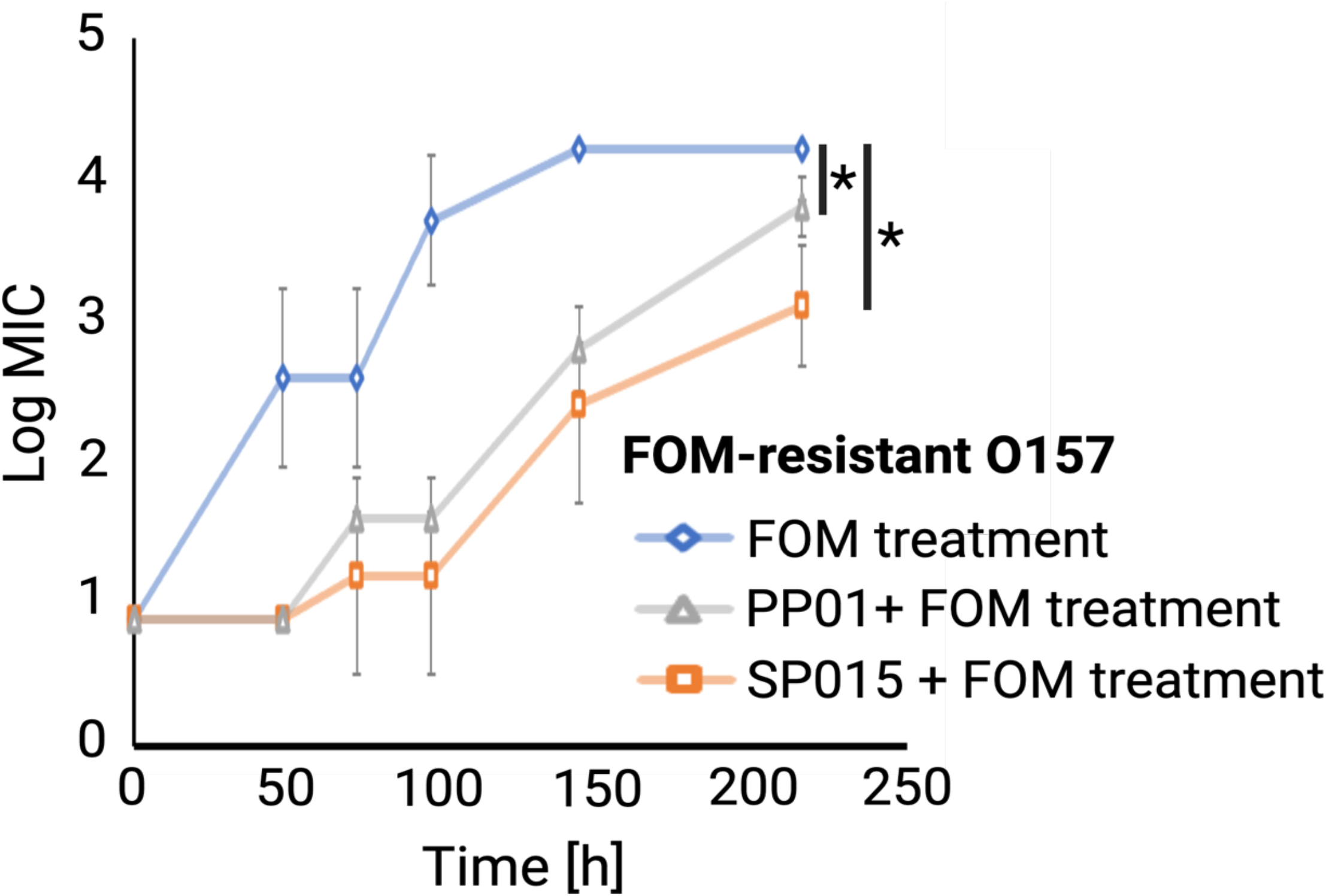
The MIC value of FOM of bacteria isolated from each round of FOM treatment (blue diamond), co-culture with PP01 and FOM treatment (grey triangle), and co-culture with SP015 and FOM treatment (orange square). Statistical difference (*P* < 0.05) is indicated by Asteric.

### The combination of phage and antibiotic delayed the emergence of phage-resistant O157

In the co-culture of bacteria and phages, a coevolutionary arm-race often occurs, whereby the bacteria generally alter the phage receptor to avoid phage adsorption, whereas the phage could evolve a mutation in the receptor-binding protein that enables the phage to use an alternative receptor (35, 40–42). A similar phenomenon might occur in the case of combination treatment with phage antibiotics; thus, to evaluate the dynamics of phage-bacteria interactions in this study, we isolated the spontaneous mutant phage and bacteria from every round of co-culture and subjected them to bacteriophage spot assays (Figs. 4A and 4 B).

**Figure 4.**
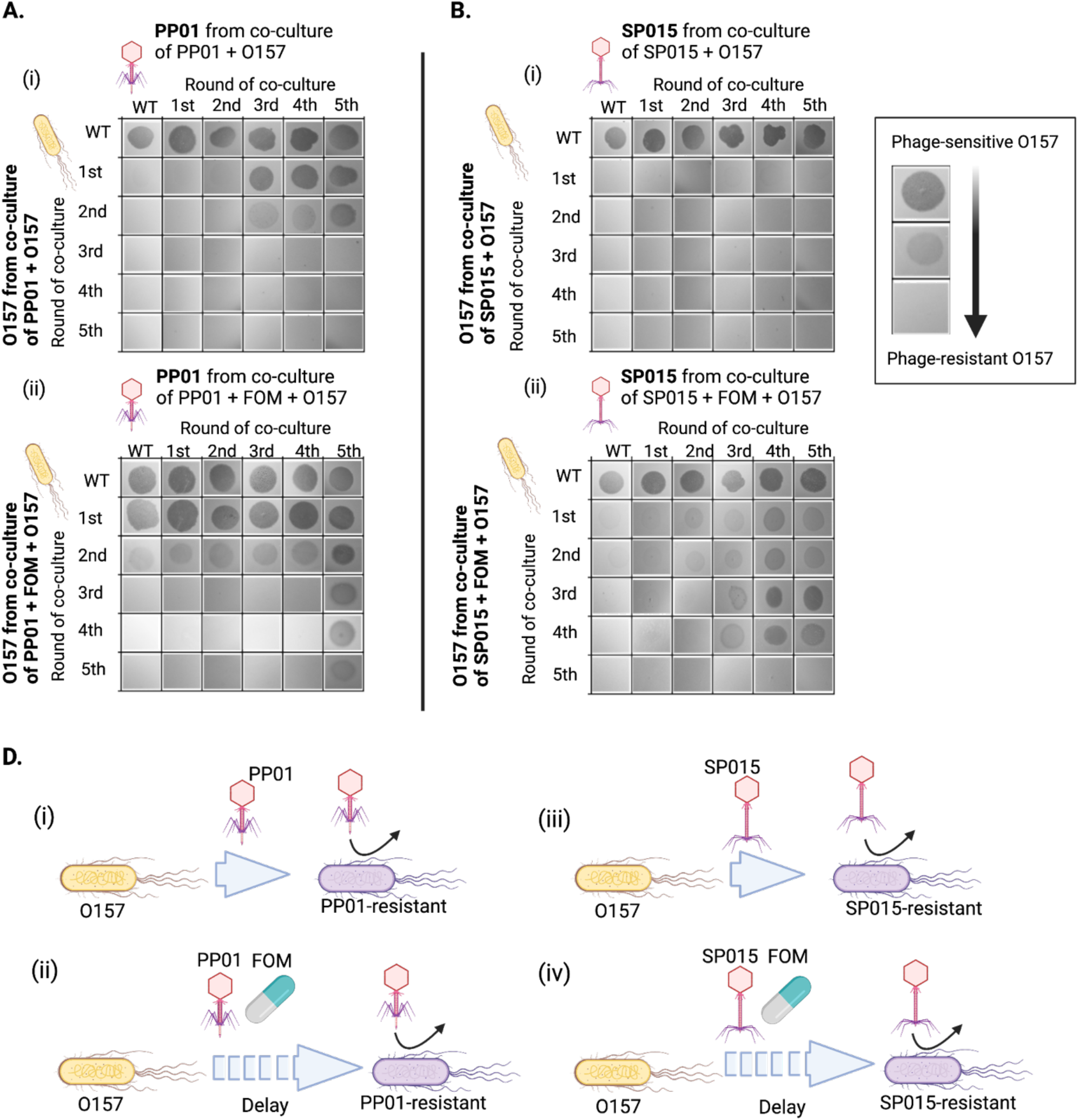
Combination of phage and FOM delay the emergence of phage-resistant clone. (A) Increased resistance of mutant O157 and counter adaptation of mutant phage PP01 in the co-culture without FOM (i) and with FOM (ii). In the absence of FOM, complete resistance of O157 against all isolated mutant PP01 was observed on resistant O157 clone isolated from third until fifth round of co-culture, whereas in the presence of FOM, no resistant clone with complete resistant was obtained until fifth round of co-culture. (B) Increased resistance of mutant O157 and counter adaptation of mutant phage SP015 in the co-culture without FOM (i) and with FOM (ii). In the absence of FOM, resistant O157 clone against all isolated mutant SP015 was rapidly isolated from first until fifth rounds of co-culture, whereas in the presence of FOM, resistant clone with complete resistant was obtained from the fifth round. (C) Schematic illustration of the emergence of phage-resistant clone in the presence or absence of FOM.

In the co-culture where FOM was absent (Fig. 4A[i]), wild-type PP01 was not able to infect O157 isolated from all rounds of co-culture, indicating that the bacteria developed resistance throughout the co-culture. The mutant PP01 phage from the first and second rounds did not form plaques against O157 isolated from the first to fifth rounds, but mutant PP01 from the third, fourth, and fifth rounds showed lytic activity against these bacteria, indicating that the phage was able to adapt to PP01-resistant O157. However, the phages became non-infective against O157 from the third to fifth rounds, possibly because of an additional mutation in O157. In the case of SP015, wild-type SP015 and mutant SP015 isolated from the first and second rounds did not form plaques on the lawns of O157 isolated from the first to fifth rounds. In contrast, mutant SP015 from the third to fifth rounds showed only slight bacteriolysis ability against O157 from the first round but did not form plaques on O157 in later rounds, indicating that SP015 failed to evolve a mutation that could counteract SP015-resistant O157.

Notably, in the presence of FOM, both PP01 and SP015 remained infectious toward bacteria isolated from the first to fourth co-cultures, indicating that resistance development in bacteria was delayed (Figs. 4A[ii]) and 4B[ii]). In summary, we conclude that bacteria easily acquire mutations and gain resistance against phages (Figs. 4C[i] and 4C[iii]). Phage PP01 was able to adapt and regain infectivity against PP01-resistant bacteria; however, the bacteria shortly acquired mutations until no infectious mutant phage could evolve further. Nevertheless, in the presence of antibiotics, although phage-resistant bacteria seem inevitable, regardless of the type of phage, the development of resistance was significantly delayed (Figs. 4C[ii] and 4C[iv]).

### Gene mutations acquired in O157 co-cultured with phage and FOM

Phage-resistant O157 was isolated from the fifth round of phage and bacterial co-culture in the absence or presence of FOM and subjected to whole-genome sequencing. Comparison of the resistant strain with the wild strain O157 revealed mutations conferring resistance to phages and FOM (Supplementary Table. S4). The number of mutations identified in resistant clones is shown in Fig. 5. The highest number of mutations was observed in FOM-resistant O157, which had nine insertion/deletion (indel) mutations and 35 point-mutations. Only two point-mutations were found in either PP01 or SP015 resistant O157. In contrast to the SF-resistant strain, which exhibited four point mutations and five indel mutations, the PF-resistant strain displayed five point-mutations and two indel mutations. In PP01-resistant O157, a nonsense mutation was identified in which the 76^th^ glutamine of OmpC (locus tag: FNZ21_13245), the receptor of PP01(39, 43), was replaced by a stop codon. In addition, arginine at position 143 of the glycosyltransferase (FNZ21_14180), an enzyme involved in the biosynthesis of oligosaccharides and polysaccharides, was replaced with a stop codon. In SP015-resistant O157, a nonsense mutation was identified, in which tryptophan at position 511 of FhuA (FNZ21_00755) was replaced by a stop codon. In addition, proline at position 696 of the DEAD/DEAH box helicase (FNZ21_02015), which is involved in various aspects of RNA metabolism, was replaced by leucine. In FOM-resistant O157, mutations were found in two transporters, UhpT (FNZ21_06865) and GlpT (FNZ21_13145), which take up FOM into the bacteria. A single base deletion of glycine at position 141 in UhpT caused a frameshift, and the amino acid at position 202 was replaced with a stop codon. In addition, the glycine at position 358 was replaced by serine in GlpT. In PF (PP01 + FOM)-resistant O157, mutations in UhpT and GlpT were also observed; serine at position 5 was replaced by a stop codon in UhpT and 555 bp from the stop codon was deleted in GlpT. In contrast, no mutations were found in OmpC, the PP01 receptor. In SF (SP015 + FOM)-resistant O157, there were no mutations in UhpT, but 57 bp was deleted within the gene encoding UhpA (FNZ21_06880), the activator of UhpT(44, 45). In addition, the aspartic acid at position 88 of GlpT was replaced with glutamic acid. Moreover, there were two mutations in FhuA, with a substitution of aspartic acid at position 218 for asparagine and a 69 bp deletion.

**Figure 5.**
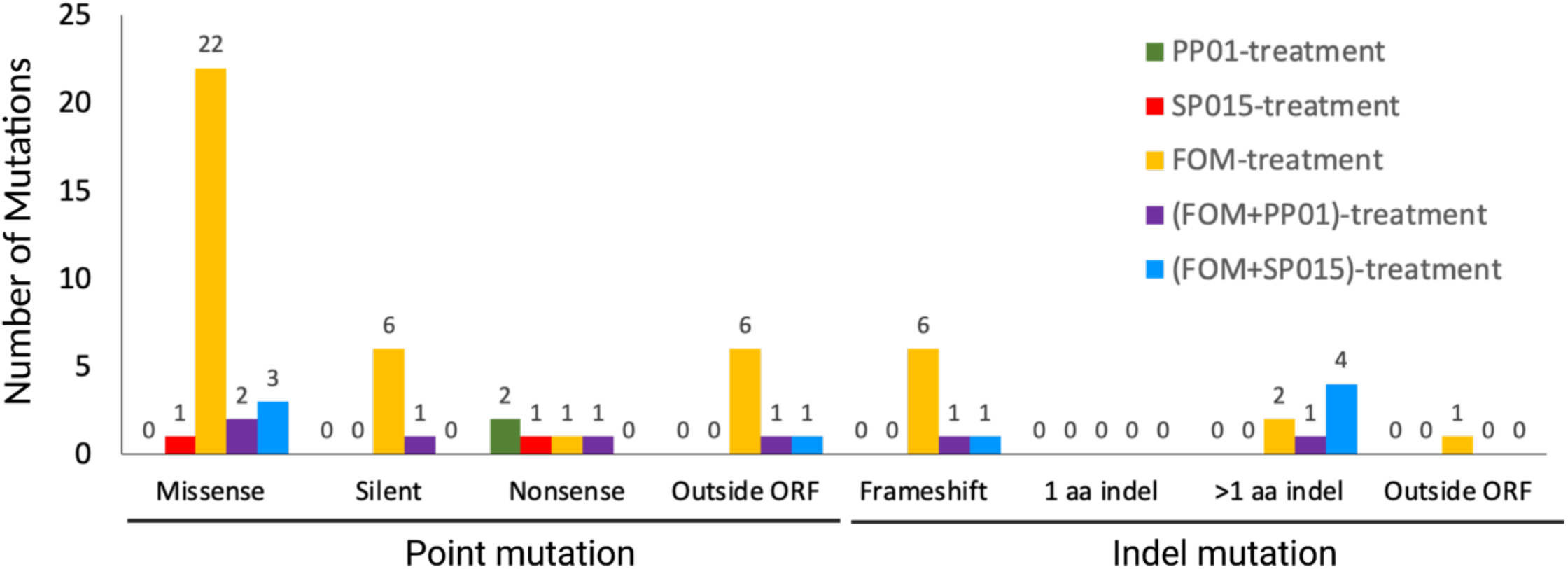
Number of mutations in the resistant bacteria isolated from fifth round of co-cultures.

Similarly, the whole genome sequence of the mutant phage was compared with that of the wild-type phage (Supplementary Table. S4). Five mutations were found in mutant PP01 from PP01 co-cultured with O157, excluding the silent mutation, including a frameshift and deletion of the termination codon due to a single nucleotide loss of leucine at position 105 of gp33, which binds to RNA polymerase and is required for the transcription of late genes such as the phage outgrowth protein. In addition, two mutations were found in the gene encoding gp37, the long-tail fiber of the phage, with a substitution of aspartic acid at position 650 for asparagine and a deletion of 687 bp. Two missense mutations were found in mutant SP015 from SP015 co-cultured with O157, both in the gene encoding the tail fiber, whereas a 7698 bp deletion was found in mutant SP015 isolated from SF co-culture.

### Identification of genetic determinant responsible for the resistance phenotype against phage and FOM

To determine the mutations responsible for FOM and phage resistance in O157, we constructed various deletion mutants of O157 based on mutations identified in the resistant clone from the co-culture (Fig. 6). We searched for candidate genes for the deletion construct by comparing the mutated genes identified in resistant clones that resist phage alone or both phage and FOM (Fig. 6A). To create the deletion mutants, four genes encoding UhpT, GlpT, OmpC, and FhuA were chosen. Since the effect of deleting the gene encoding UhpA (the activator of UhpT) should be the same as that of deleting the gene encoding UhpT(44, 45), this gene was not selected for deletion. As shown in Fig. 6B, we confirmed that PP01 and SP015 could not form plaques nor adsorb onto the OmpC and FhuA deletion mutants, respectively, suggesting that OmpC is a receptor of PP01, whereas FhuA is a receptor of SP015. While OmpC has been previously reported as a receptor for PP01 (39, 43), in the current study, we found that SP015 could not adsorb onto FhuA deletion mutant O157, and complementation with FhuA restored the adsorption; thus, we confirmed that FhuA is a receptor for SP015 (Supplementary Fig. S3D). Notably, while wild-type PP01 requires the OmpC protein as a receptor, mutant PP01 from the fifth round of PP01 co-cultured with O157 could infect OmpC-null O157, indicating that mutant PP01 could utilize different receptors for infection. Deletion of OmpC did not change the MIC values, whereas FhuA deletion mutants showed lower MIC values than wild-type O157. The strains lacking UhpT and GlpT did not differ from the wild-type in their sensitivity to phages. While the MIC values of wild-type O157 were 16 μg/mL, those of mutants *ΔuhpT* and *ΔglpT* were 32 and 16 μg/mL, respectively. Double deletion of UhpT and GlpT, but not a single deletion of the gene, causes O157 to become FOM-resistant (MIC ζ 256 μg/mL) while retaining the bacteria susceptible to phages. Remarkably, triple deletion of OmpC, UhpT, and GlpT *(ΔUhpTΔGlpTΔOmpC*) resulted in a two-fold increase in theMIC value compared to the UhpT and GlpT double deletion *(ΔUhpTΔGlpT*) mutant. In contrast, the MIC of *ΔUhpTΔGlpTΔFhuA* was the same as that of *ΔUhpTΔGlpT*. Thus, our findings highlight a possible role of OmpC in FOM uptake by bacteria.

**Figure 6.**
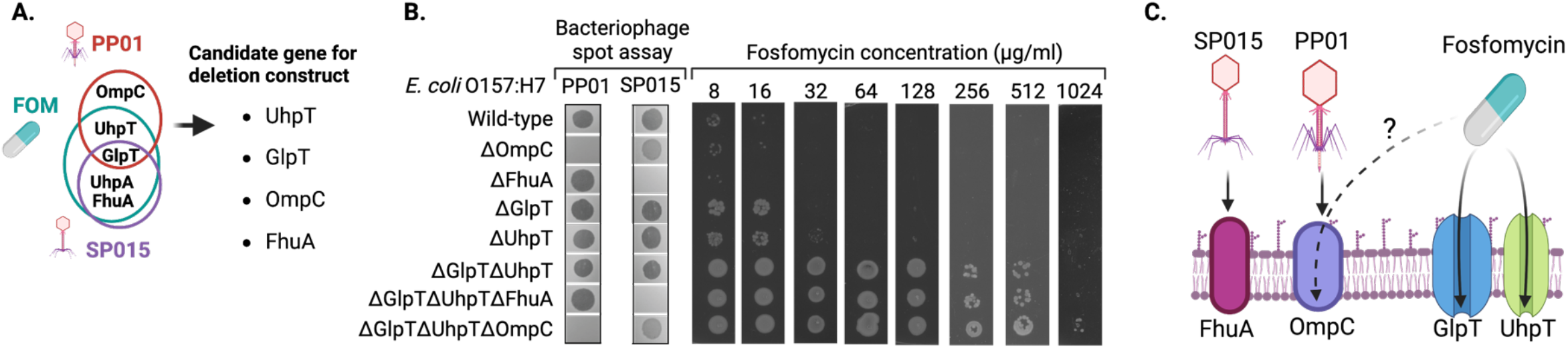
Common mutation identified in resistant bacteria from FOM treatment, phage alone, or phage with FOM treatment. (A) When phage and antibiotics were added to O157, five genes were mutated in common. Since the promoter activity of *uhpT* is absolutely dependent on UhpA, four genes (*uhpT, GlpT, ompC* and *fhuA*) were selected for construction of deletion mutant. (B) The sensitivity to phage and MIC values of FOM changed in the deletion mutant. In O157 strains lacking OmpC and FhuA proteins, PP01 and SP015 phage infection was inhibited, respectively. O157 becomes FOM-resistant when both UhpT and GlpT proteins are deleted. Triple deletions of UhpT, GlpT and OmpC increase the MIC by twofold compared to the double deletion mutant of UhpT and GlpT. (C) Schematic representation of the phage receptor and FOM uptake channel observed in this work. SP015 and PP01 likely recognize FhuA and OmpC as their respective receptors. FOM enters cell through GlpT and UhpT, and additionally, it may be able to enter the cell through OmpC.

In conclusion, we found that the phages used in this study exploited different receptors: PP01 utilized OmpC, whereas SP015 recognized FhuA (Fig. 6C). Deletion of either GlpT or UhpT did not significantly increase the MIC value of FOM (Fig. 6B), but deletion of both significantly increased the MIC value, indicating that FOM enters bacteria through at least two different receptors, GlpT and UhpT (Fig. 6D). The absence of OmpC and two FOM channels increased the MIC more than it did with the deletion of two FOM channels, indicating that FOM may be able to enter the cell through OmpC (Fig. 6E).

## Discussion

The combination of phages and antibiotics has been widely used to enhance the eradication of drug-resistant bacteria and mitigate the spread of antibiotic resistance worldwide (8, 46, 47). Several studies have examined the mechanisms underlying phage-antibiotic synergy. Owing to the selection pressure on bacteria caused by phage infection, some toxicity, drug sensitivity, and growth factors are lost, and phage-resistant strains are often less toxic, more sensitive to antibiotics, and grow at a slower rate than wild-type strains (46, 47). Combining phages with antibiotics can reduce resistant clones, as our study showed, but the type of phage utilized will affect the outcome (Fig. 1C-1G). The best inhibition of the resistant clone development was shown by the combination of FOM and SP015. Although not significant, the fitness cost caused by the combination treatment was observed in the current study (Supplementary Fig. S6).

Phage adsorption is the first critical step for phages to infect bacterial hosts; therefore, mutations in the receptor are commonly found in bacteria as a potent defence strategy to escape phage predation (12, 40, 48–50). The appearance of a mutant phage capable of infecting some phage-resistant bacteria was observed after five rounds of co-culture (Fig. 4A (i, iii, and iv)) and was speculated to be associated with the mutation found in the tail protein.

In the culture under FOM without phage addition, the MIC was measured at every end of the rounds, and an MIC exceeding the upper limit of 16,384 μg/mL was observed in all five runs within three rounds of co-culture (Fig. 3). Although FOM has been reported to be effective in eradicating O157 (25), the rapid emergence of FOM-resistant O157, which has an MIC 1,024 times higher than that observed in O157 wild-type, indicate that O157 can easily develop resistance against FOM. Whole-genome analysis of FOM-resistant O157 revealed mutations in the genes encoding the two transporters (UhpT and GlpT) that take up FOM. It is believed that the permeability of FOM is reduced and the level of resistance is increased by the accumulation of mutations, including numerous additional genes found in the FOM resistant clone (Supplementary Table S4), but the mechanism behind these changes is yet unknown.

In the co-culture in which phages and antibiotics were added (PP01+FOM, SP15+FOM), a synergistic effect of the combination was observed in the first round, and the emergence of resistant bacteria was suppressed more than when each phage was used alone. In particular, the combination of SP015 and FOM significantly suppressed the emergence of resistant bacteria compared to the combination of PP01 and FOM, suggesting that an optimal combination of phages and antibiotics should be achieved in antimicrobial therapy. Throughout the five rounds of co-cultures, the average MIC values of resistant O157 from PP01+FOM and SP015+FOM were significantly lower than those of resistant O157 from FOM treatment alone (Fig. 3), suggesting that the combined use of phage and antibiotics suppressed the level of resistance to antibiotics. Furthermore, mutant phages infected with resistant O157 were continuously available in the co-culture PP01+FOM or SP015+FOM until the last round of co-culture (Fig. 2A (ii) and 2 B (ii)), suggesting that the phage and antibiotic combination could suppress the emergence of phage-resistant bacteria in addition to the emergence of antibiotic resistance.

Whole-genome analysis of the strains resistant to PP01+FOM showed that mutations occurred in the same genes (*uhpT* and *glpT*) as those in FOM-resistant O157. In contrast, no mutations were found in o*mpC*, the PP01 receptor gene, but point-mutations were found in *hldE* (FNZ21_08945), a gene involved in LPS biosynthesis. The adsorption of PP01 onto O157 is divided into two stages: first, the long tail fiber (gp38) reversibly attaches to the host receptor (OmpC), and then the short tail fiber (gp12) irreversibly attaches to LPS (32, 39, 43). Therefore, it is possible that a mutation in *hldE* prevented PP01 from adsorbing onto the host. Whole-genome analysis of SP015+ FOM-resistant O517 showed mutations in the two transporters for FOM uptake and in the receptor of SP015 (FhuA). These results suggest that O157 evolves mutations in the phage receptor and FOM uptake channel to escape phage and FOM suppression.

To further clarify the mutations observed in resistant O157, we constructed various deletion mutants of O157. Deletion of the phage receptor gene resulted in the loss of plaque-forming ability of each phage. This is thought to be due to the inability of phage ligands to attach to host receptors. Strains lacking one transporter (UhpT or GlpT) did not show a significant increase in the MIC of FOM compared to the wild-type, but when both were deleted, the MIC was significantly increased, indicating that FOM could enter the cell when one of the transporters was available. Cases of FOM resistance owing to reduced permeability caused by mutations in this transporter have been reported in a previous study (25). In this experiment, we created a strain deficient in both genes for the phage receptor and FOM uptake channel. The phage sensitivity of *ΔUhpTΔGlpTΔOmpC* was the same as that of *ΔOmpC*, but the MICs values of FOM in *ΔOmpC* and *ΔUhpTΔGlpT* were 16 and 512 μg/mL, respectively, whereas *ΔUhpTΔGlpTΔOmpC* was 1024 μg/mL. This may be due to the lack of two transporters, which reduces FOM permeability into the bacteria; however, because of the small molecular weight of FOM, it is transmitted through other transporters. In the present case, OmpC corresponded to this, and it is thought that the MIC value increased because *ΔUhpTΔGlpTΔOmpC* is also deficient in OmpC. Our study highlighted the potential use of phage and antibiotic treatment to control O157; however, we should be more selective in determining which type of phage should be used to achieve the best outcome.

In nature, phages impact bacterial evolution through an antagonistic arms race with bacteria. Therefore, unlike antibiotics, phages often counter-adapt and overcome phage resistance in bacteria (40, 48). Current study showed that phages counter-adapt to phage resistance even when phage and FOM are used together (Fig. 4A(i)). Since phage-resistant bacteria acquired mutations in genes encoding membrane proteins and lipopolysaccharides, both of which have been reported as virulence factors in Gram-negative bacteria, combination therapy of antimicrobials and phages against O157 may be applicable to other antimicrobials other than FOS (Supplementary Table S4) (40, 51–57). However, our study was limited to *in vitro* analysis, additional observation of *in vivo* experiments is necessary to have more comprehensive understanding on the potential use of combined phage antibiotic therapy prior to clinical application. Careful selection of phages is important and precise phage-antibiotic combinations should be examined to see whether they work synergistically to combat bacterial infection. Furthermore, while our current study is limited to a single phage application, we believe that combining numerous phages as a cocktail with antibiotics may considerably decrease resistant bacteria and boost their therapeutic efficacy.

## Acknowledgement

The figure in this article was created using BioRender.com (accessed August 2022).

---

**Supplementary Table S1.** Bacteria, Bacteriophage, and plasmid used in this study

| Strain name | Description |
| --- | --- |
| **Bacteria** |  |
| *E. coli* O157:H7 ATCC43888 | Wild-type (no shiga toxins 1 and 2) |
| R5-PP01 | PP01-resistant O157:H7 (Round 5th) |
| R5-SP15 | SP015-resistant O157:H7 (Round 5th) |
| R5-FOM | Fosfomycin-resistant O157:H7 (Round 5th) |
| R5-PF | PP01+FOM-resistant O157:H7 (Round 5th) |
| R5-SF | SP015+FOM-resistant O157:H7 (Round 5th) |
| *E. coli* O157:H7 EDL933 |  |
| *E. coli* O157:H7 CR-3 |  |
| *E. coli* O116:H10 ATCC23541 |  |
| *E. coli* O157:H19 A2 |  |
| *E. coli* O157:H37 CE273 |  |
| *E. coli* K12 JM109 |  |
| *E. coli* K12 JM110 |  |
| *E. coli* K12 W3110 |  |
| *E. coli* C |  |
| *E. coli* C600 |  |
| *E. coli* BL21 |  |
| *E. coli* DH10B |  |
| *E. coli* DH5α | |
| *E. coli* MC101 |  |
| *E. coli* Top10 |  |
| *E. coli* HST08 |  |
| *E. coli* O157:H7 ΔuhpT | *uhpT* knockout O157 |
| *E. coli* O157:H7 ΔglpT | *glpT* knockout O157 |
| *E. coli* O157:H7 ΔuhpTΔglpT | *uhpT* and *glpT* O157 |
| *E. coli* O157:H7 ΔompC | *ompC* knockout O157 |
| *E. coli* O157:H7 ΔfhuA | *fhuA* knockout O157 |
| *E. coli* O157:H7 ΔuhpTΔglpTΔompC | *uhpT*, *glpT* and *ompC* O157 |
| *E. coli* O157:H7 ΔuhpTΔglpTΔfhuA | *uhpT*, *glpT* and *fhuA* O157 |
| **Phage** |  |
| PP01 | Wild type, infectious to O157:H7 |
| SP15 | Wild type, infectious to O157:H7 |
| m5-PP01 | mutant PP01 in the PP01 condition (Round 5th) |
| m5-SP15 | mutant SP15 in the SP15 condition (Round 5th) |
| m5-PF | mutant PP01 in the PP01+FOM condition (Round 5th) |
| m5-SF | mutant SP15 in the SP15+FOM condition (Round 5th) |
| **Plasmid** |  |
| pKOV | Temperature-sensitive vector; Cm<sup>r</sup> *sacB* |
| pK-uhpT | For knockout of O157 *uhpT* |
| pK-glpT | For knockout of O157 *glpT* |
| pK-ompC | For knockout of O157 *ompC* |
| pK-fhuA | For knockout of O157 *fhuA* |
| pFhuA | For complementation of *fhuA* |

**Supplementary Table S2.**
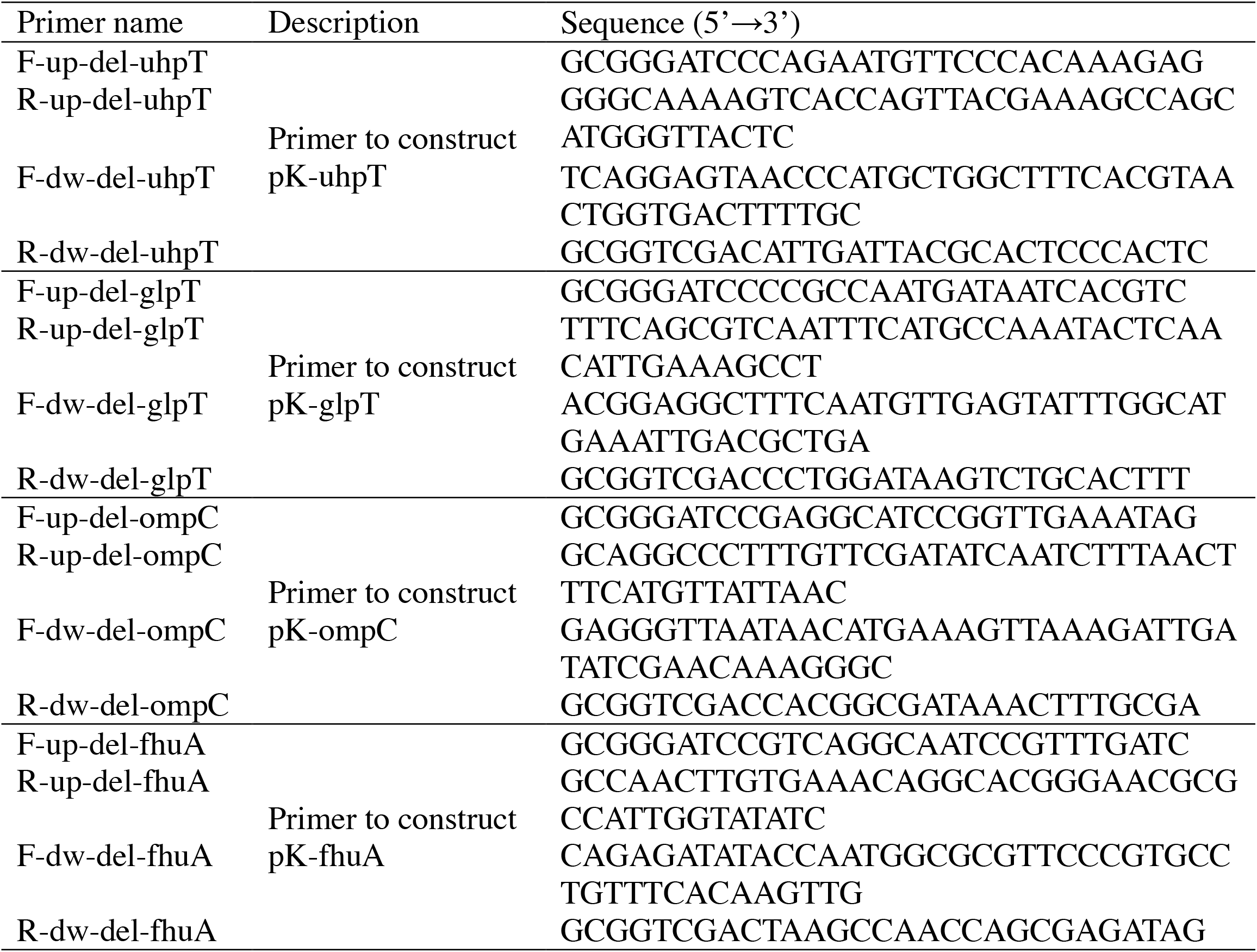
Primers used in this study.

**Supplementary Fig. S3.**
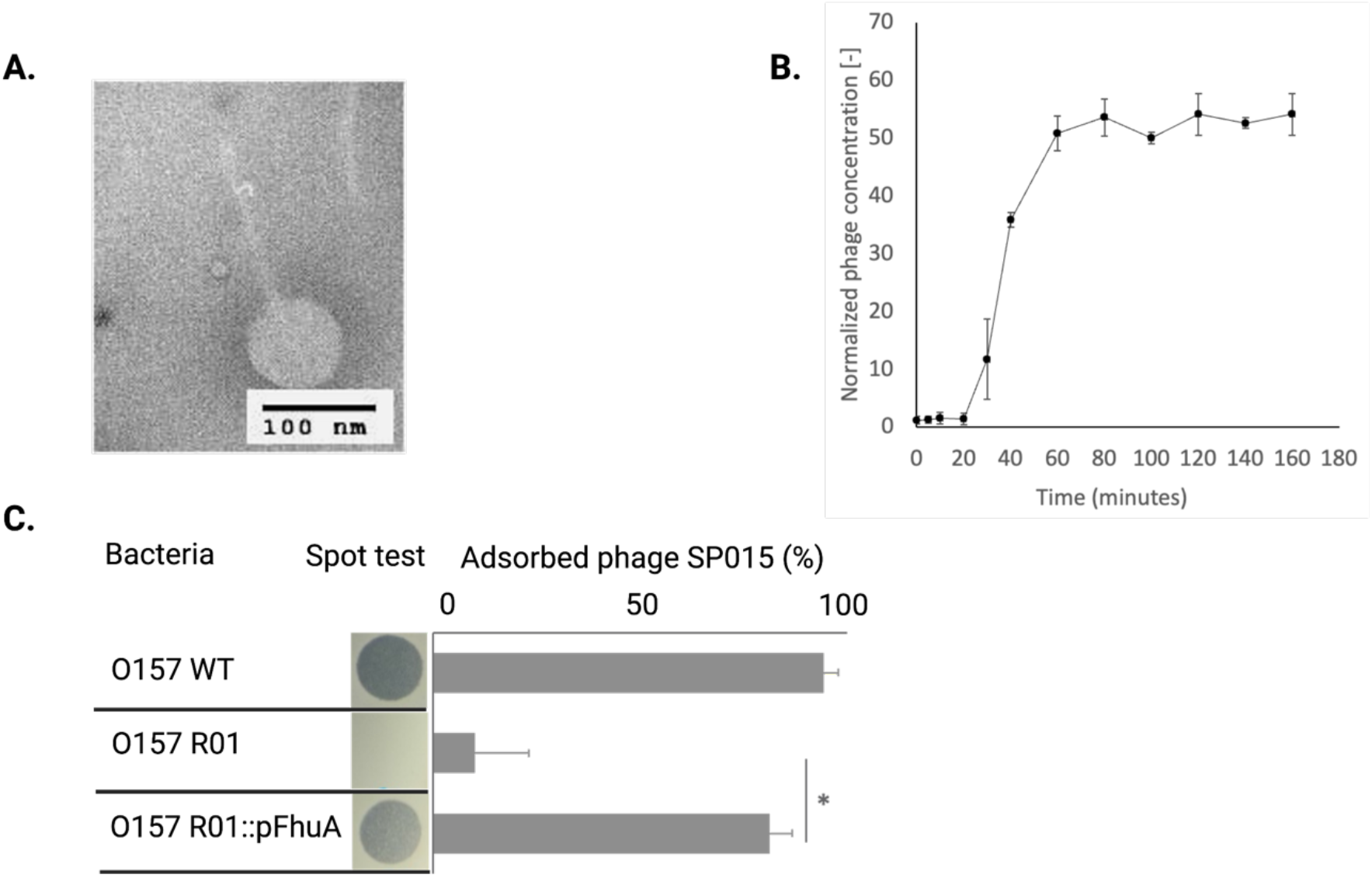
Characterization of SP015. (A) Morphological observation of SP015 under TEM. (B) One-step growth of SP015. (C) Receptor identification by using spontaneous mutant bacteria with truncated FhuA.

**Supplementary Table. S4.**
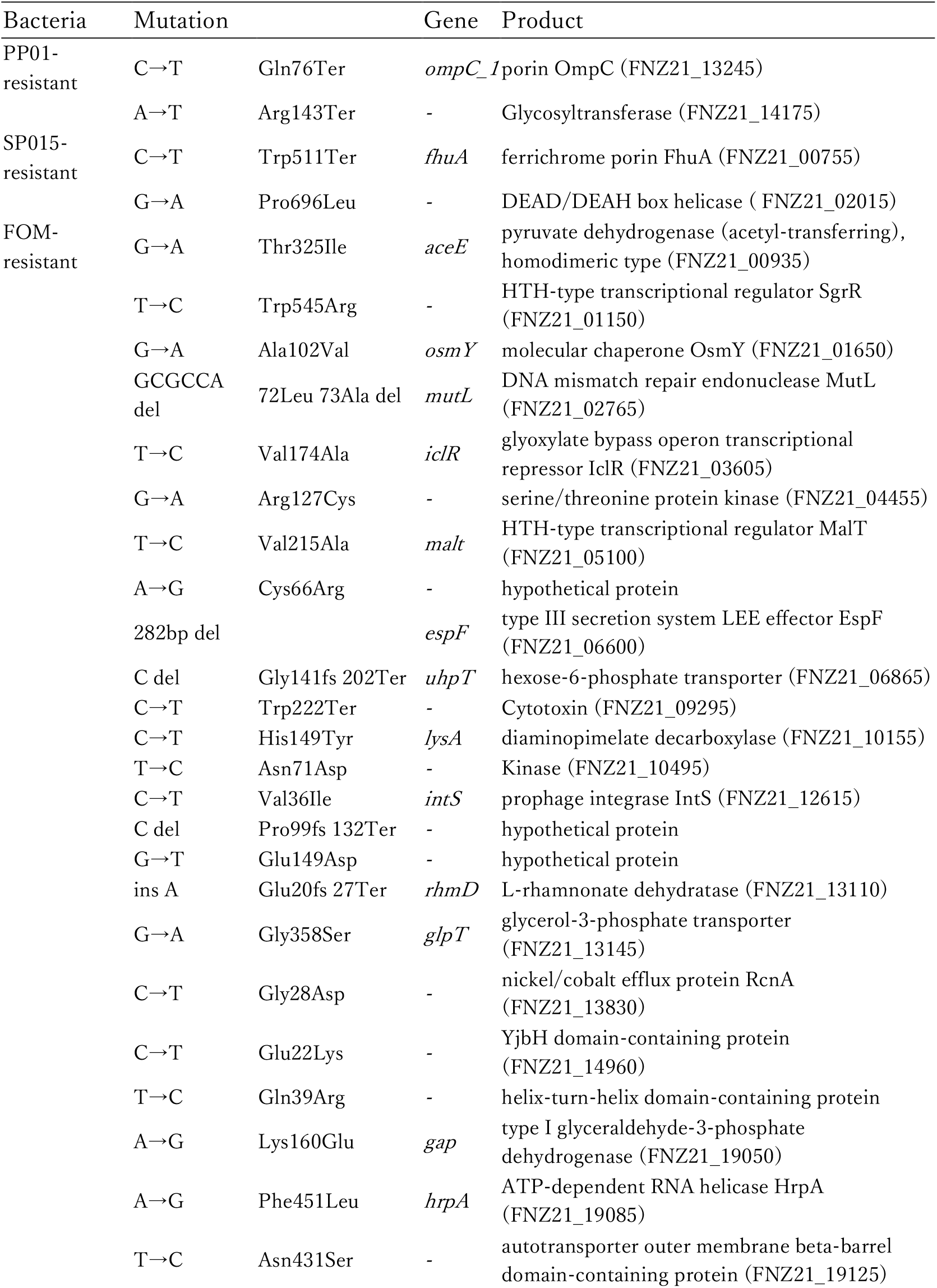
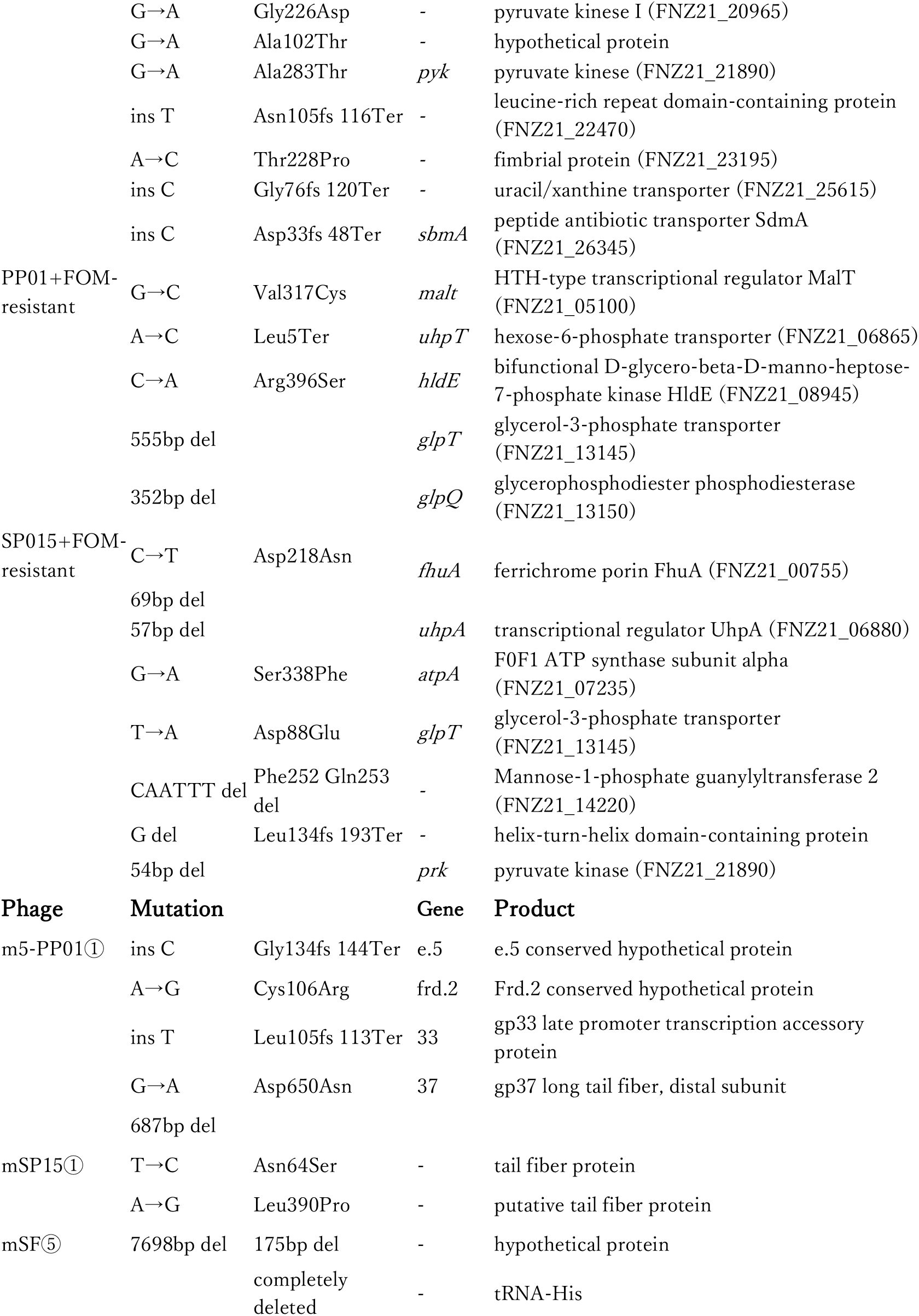
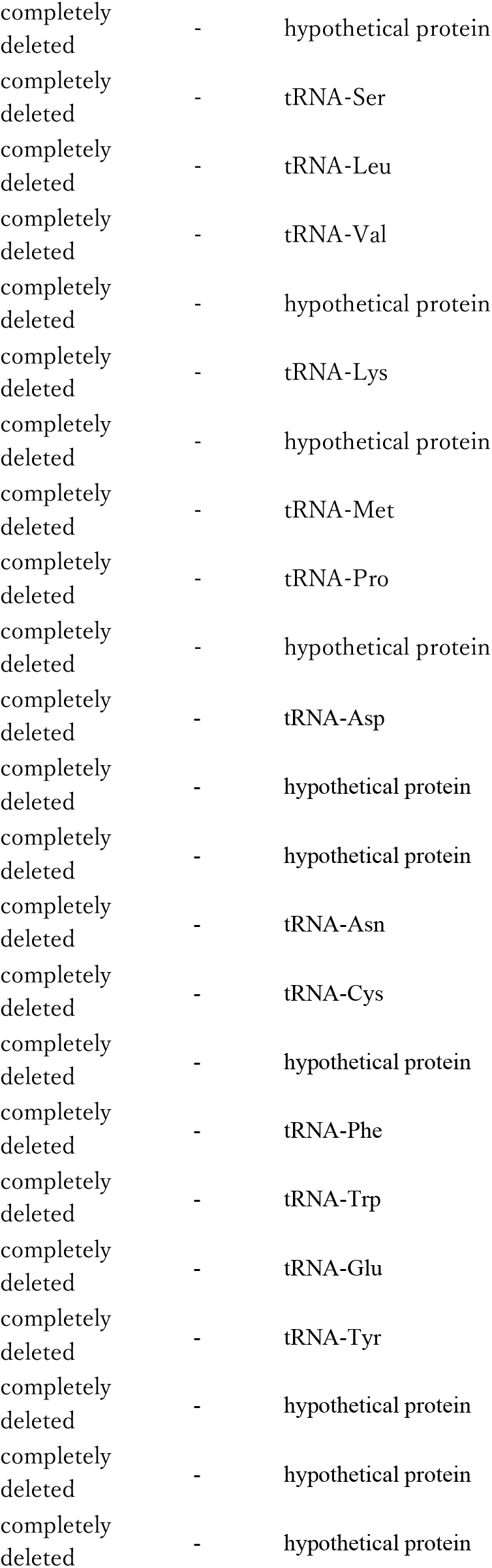
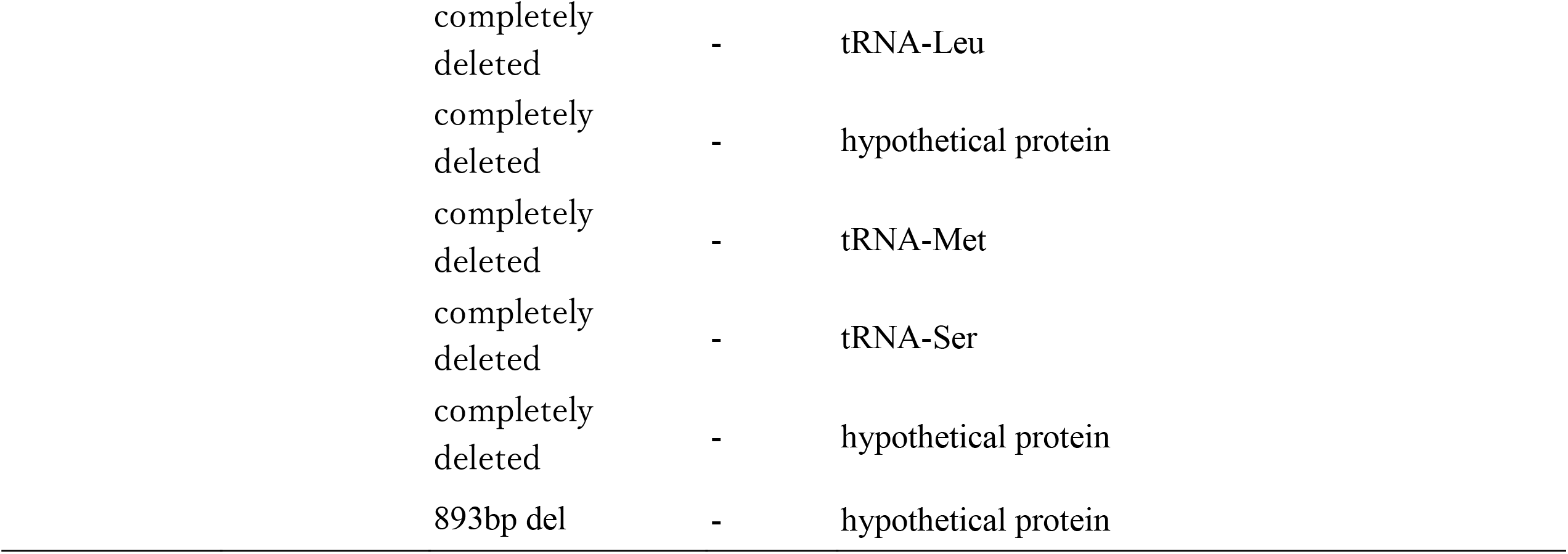
Mutation identified in resistant bacteria and mutant phages.

**Supplementary Fig. S5.**
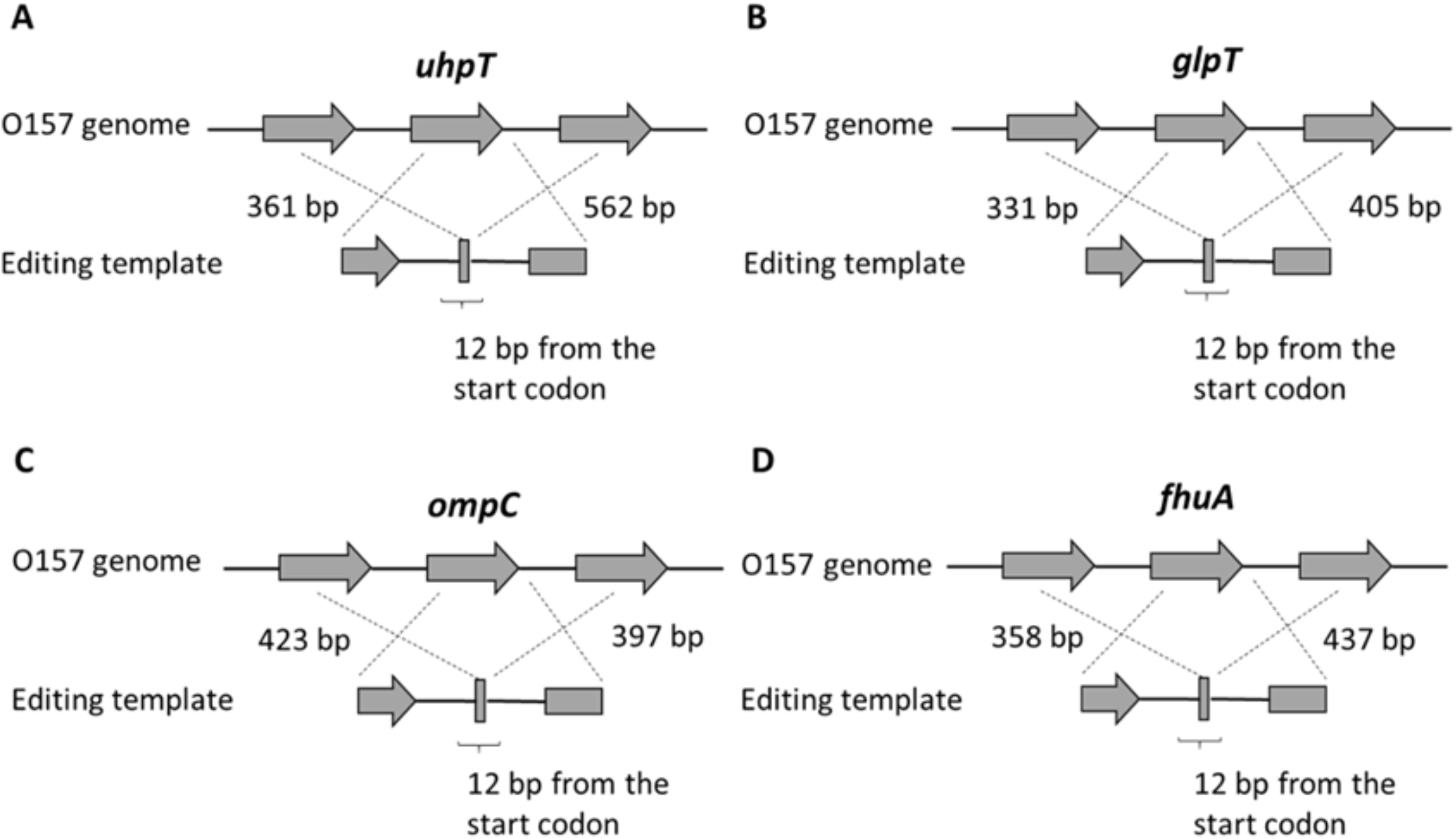
Plasmid construction to delete gene of interest.

**Supplementary Fig. S6.**
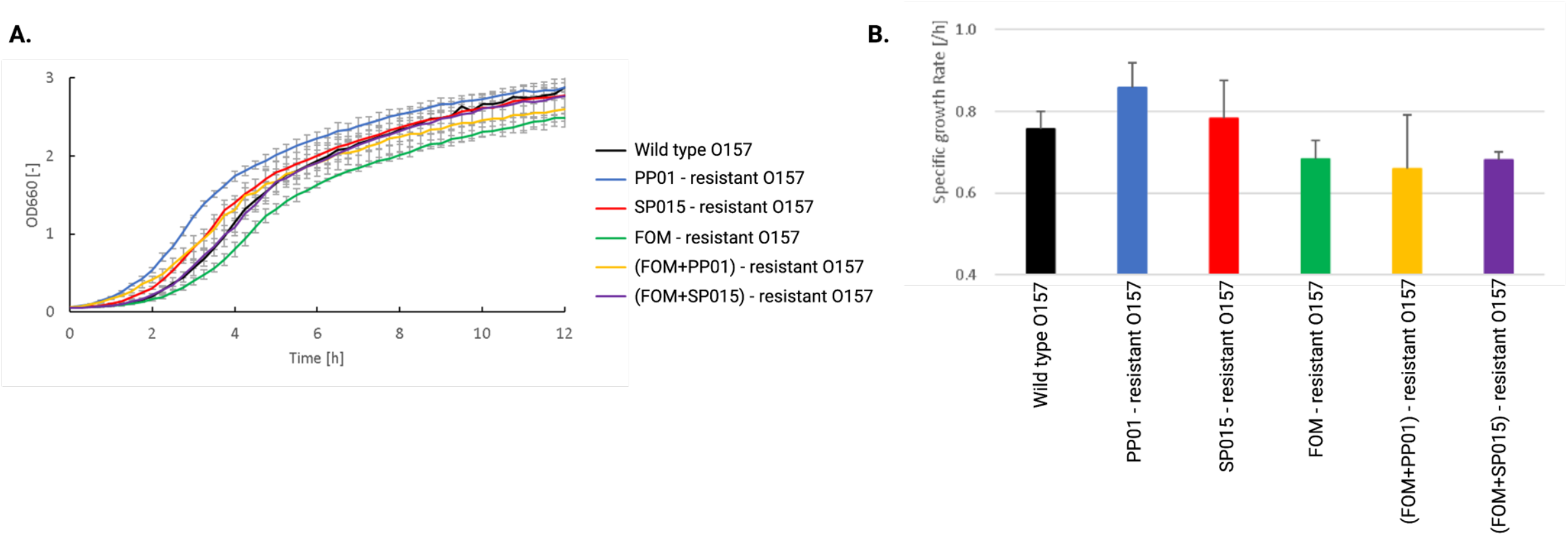
Fitness cost in resistant O157. (A) Bacterial growth in liquid medium. (B) Specific growth of bacteria extracted from (A). The experiment was conducted in three biological triplicates.

